# Human body prediction of size and shape: a hormonal framework

**DOI:** 10.1101/722777

**Authors:** Jeroen van Vugt

**Affiliations:** Krossfields, Vught, 5262AK, The Netherlands

## Abstract

To achieve high prediction accuracy with minimal inputs from online retail respondents, a method was developed and tested to predict the size and shape of the human body in 3D using a hormonal framework. The prediction method is based on geometric morphometrics, image analysis, and kernel partial least squares regression. The inputs required are answers to three closed-ended questions and a passport photo. Prediction accuracy was tested with the 3D body scan dataset of the Civilian American and European Surface Anthropometry Resource project. Results from the test dataset showed that approximately 82% of the error expectations of landmarks followed a log-normal distribution with an expectation of 8.816 mm and standard deviation of 1.180 mm. The remaining 18% of the error expectations of landmarks followed a log-normal distribution with an expectation of 18.454 mm and standard deviation of 8.844 mm, which may herald future research. Benchmarked with another method, the proposed method features much less input. In addition to high accuracy, the method in this paper allows for visualisation of results as real-size meshes in millimeters.

## Introduction

Hormones may be described as chemical signals excreted into blood vessels by cells, and these signals have a coordinating action via receptor binding particularly on distant target cells^1^. The target cell response, which is contingent on hormonal presence, sensitivity, and cell programming, can lead to survival or apoptosis (programmed cell death), differentiation, proliferation, and/or execution of a specialised function, such as contraction or secretion^2^.

The effect of hormones on the size and shape of the human body may be illustrated when their levels drop significantly, as can be the case for schizophrenia. This condition may be associated with a narrowing and reduction of the lower and middle facial region, the forehead and nasal bone^3^, as well as body linearity and low Body Mass Index (BMI)^4^. The cause of the condition may be a deficit of the hormone oestrogen^5^, which is important in bone metabolism^6^.

Eight categories of hormones can be highly relevant to human morphology, namely: (1) growth^7,8,9^, (2) oestrogen^10^, (3) testosterone^11^, (4) dihydrotestosterone^12^, (5) metabolism and food intake^13,14^, (6) bone^15,16^, (7) glucocorticoid^17^, and (8) breastmilk secretion^18^. The impact of these categories may differ between sexes^19^, life phases^20,21,22,23^ (childhood, puberty, and adulthood), populations^24^, as well as change during pregnancy^18^. In this article, population refers to a group of humans that is morphologically distinct due to development and evolution.

A hormonal framework has been used in this article to create a prediction method of the size and shape of a human body in 3D, and to investigate its prediction accuracy. To the knowledge of the author, no other prediction methods published in the scientific literature have been informed by hormonal theory. To predict human body shape, Zhu and Mok^25^ created a method with two orthogonal-view photos of the whole dressed body as input. The method’s disadvantages are that it requires much input and prediction of the body shape is dependent upon the fit of clothing. Xi, Guo, and Shu^26^ used predictive clustering trees with demographic variables to predict human body shape. However, they did not explain their methodology with a theoretical framework. Demographic variables may explain group features, but analysis has to be on the level of the individual to predict individual human body features. An individual can be predicted with a hormonal framework because hormones form the whole body. Specifically, by taking just one picture of the face, the body can be predicted. This insight is used for the first time in this article to create a prediction method that can enable online clothing stores to reduce the number of returns from clients.

With the method developed in this article, clients only have to provide as input their stature, body weight, age, and a passport photo. The method was based on a dataset of 3D body scans from the Civilian American and European Surface Anthropometry Resource (CAESAR) project^27^, geometric morphometrics, image analysis, and kernel partial least squares regression.

## Methods

The CAESAR dataset contains raw data that requires a lot of processing before they can be applied in a prediction. This section first outlines how the raw data was transformed into predicted and predictor variabeles. It then explains the statistical prediction model. The section further shows how the test dataset was selected and error measures were calculated. Finally, the benchmark prediction method was sketched.

### Variables

The dataset of the CAESAR project contains coordinates of 73 fixed landmarks per subject, rgb colour values per vertex, demographic variables, and traditional-style measurement variables such as stature, body weight, and skinfold.

Each 3D body scan was reconstructed prior to the variables being obtained. The reconstruction was carried out using a triangulation of the point cloud in mm where holes were filled using Version 2017.0.0 of computer program Geomagic Control X^[28]^. Size and shape variables were obtained with Version 3.4.4 of analysis software R^[29]^ and Version 3.0.6 of software package geomorph^30^.

Strongly asymmetric landmark configurations due to posture and/or body characteristics were removed using a Procrustes analysis with fixed landmarks. A partial generalised Procrustes analysis was carried out using function gpagen; to center, scale to a centroid size of 1, and rotate all configurations^31^. The partial Procrustes distance *d_P_* was used, the square root of the sum of squared differences between homologous points^32^. Size of a configuration, for example of configuration *M*, was denoted by centroid size:

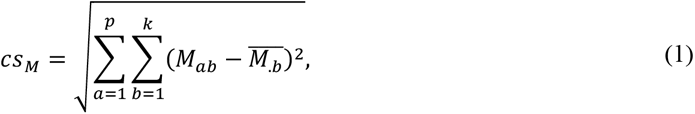

where, along with *p* landmarks, the *k* centroid coordinates were each an arithmetic average of the coordinates of a dimension^32^. Strongly asymmetric configurations were identified by calculating the partial Procrustes distance between every subject’s shape, aligned by principal axes, and its reflection.

#### Predicted variables

Additional landmarks were required to predict as many body characteristics as possible. A nose tip vertex with the longest Euclidean distance to the nuchale landmark was selected as fixed landmark. The nose tip landmark was added, as nose size can be proportional to nasal cavity size to allow oxygen intake. The size of the human nasal cavity can be positively related to lean body mass, since more tissue maintenance requires more oxygen consumption^33^. In addition to this, function buildtemplate was used to place surface sliding semilandmarks on a mesh of a subject following the method of Gunz, Mitteroecker, and Bookstein^34^. Those semilandmarks were projected to the mesh of every other subject using function digitsurface.

An additional partial generalised Procrustes analysis was carried out with 74 fixed landmarks and 2000 surface sliding semilandmarks, followed by a selection to obtain the landmarks that best delineate the body. The shape data was aligned by principal axes, and the semilandmarks were slided with the Procrustes distance. The landmark selection was obtained by first performing k-means clustering on the average shape mesh with function fastKmeans of Version 2.6 of software package Morpho^35^. This mesh was constructed by calculating the average shape configuration using function mshape. Function findMeanSpec was used to choose the mesh of a subject whose shape was closest to the average shape. The subject’s mesh was used as input for function warpRefMesh where, using the thin-plate spline method, the mesh was transformed to the average shape. Isolated pieces of the transformed mesh were removed with function vcgIsolated of Version 0.17 of software package Rvcg^36^. Next, every fixed landmark and the Euclidean-closest landmark to each cluster center were included for further analysis. Finally, selected surface sliding semilandmarks on the hands were removed because they were misplaced when fingers were not discernable from each other.

Shape coordinates underwent an orthogonal projection in order to be able to use linear multivariate techniques in the analysis^31^:

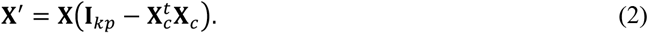

In this equation is **X** a *n × kp* matrix and **l**_*kp*_ a *kp × kp* identity matrix. Row vector **X**_*c*_ is derived from the reference, which is the average shape of which the configuration matrix *G* is scaled with its centroid size^32^:

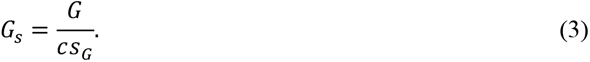

The orthogonal projection and reference were used to calculate the variables for analysis, which are Kendall tangent space coordinates^31^:

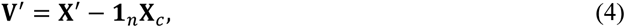

with column vector **1**_*n*_ with *n* ones. The usability of the orthogonal projection was checked by including each Euclidean distance in tangent space in a regression equation and squared correlation with corresponding Procrustes distance *ρ*^[32]^. For this, the lm and cor function were used, and the relationship between *ρ* and *d_P_* can be described as:

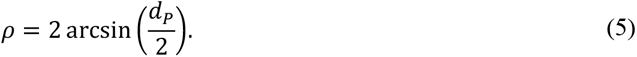

The predicted variables were the natural logarithm of centroid size and Kendall tangent space coordinates of fixed landmarks and selected semilandmarks.

Predicted Kendall tangent space coordinates were back transformed to shape configurations by approximately adding the reference^31^:

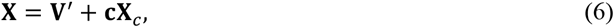

with cos (*ρ*) for each subject in column vector **c**. As an approximation for *ρ* the Euclidean distance in tangent space was used. The predicted natural logarithm of centroid size was back transformed by taking its exponent.

#### Predictor variables

The natural logarithms of configuration height in mm and body weight in kg, measured with a scale^37^, were used to predict growth hormone responses. Responses to other hormones were predicted with the shape and skin colour of the face. Oestrogen, testosterone, and hormones of metabolism and food intake can have an effect on the face shape^38,39,40^. Furthermore, face shape can be related to craniometrics, which can be used to classify humans into geographic origin^41^. Population and morphological oestrogen responses can be predicted with skin colour, which can be related to pigment melanin^42^, a decrease of which can be the result of an oestrogen reduction^43^.

The face was analysed with facial colour variables extracted from a picture of the front of the whole body mesh. First, a partial generalised Procrustes analysis was performed with fixed-landmark configurations that were aligned to principal axes. Second, onto each shape configuration its associated mesh was rotated and scaled using function rotmesh.onto. Isolated pieces of the mesh were removed using function vcgIsolated. Third, the resulting mesh was plotted without light calculation and margins using function plot3d of Version 0.99.16 of software package rgl^44^. A picture of the front of the mesh was taken with functions view3d, using a field-of-view angle of 0 degree, and rgl.snapshot. Fourth, the picture was converted to a pixmap object of Version 0.4.11 of software package pixmap^45^, devoid of white space, and cropped to obtain the face. Image cropping was performed with the landmark(s) of the cervicale, sellion, and tragion. Again, the excess white space was removed. Fifth, the resulting picture was scaled to a width and height of 30 by 30 pixels using function image_scale of Version 1.9 of software package magick^46^. This results in a picture on which hormonal responses of the subjects can still be visible, but not their age, so the natural logarithm of age in years was added to the predictor variables. Sixth, rgb variables were transformed to variables of rg chromaticity color space, making the normalised blue channel redundant. The rg chromaticity variables were used to avoid the usage of color correction algorithms that can add prediction error and require extra effort to perform. In addition, those variables are free from shading or shadow effects. Lastly, the rg chromaticity variables were transformed with the logit transformation. In order to take the natural logarithms of zero for the logit transformation, a constant of 1 was added to rgb values prior to transforming them to rg chromaticity color space.

#### Summary

Predicted and predictor variables are summarised in Table 1.

**Table 1.** Predicted and predictor variables.

|  |
| --- |
| Predicted variables: |
| • Natural logarithm of centroid size<br>• 3D Kendall tangent space coordinates of fixed landmarks and selected surface sliding semilandmarks |
| Predictor variables: |
| • Natural logarithm of configuration height in mm<br>• Natural logarithm of weight in kg<br>• Natural logarithm of age in years<br>• Logit transformations of rg chromaticity color space variables that were extracted from a face picture |
| **Table 1.** Predicted and predictor variables. |

### Statistical model

Predicted variables were included together with autoscaled predictor variables in a partial least squares regression of Version 2.7.0 of software package pls^47^, using function plsr performing kernel partial least squares regression. The amount of components to use for predictions was determined with option CV using a k-fold cross validation on the training and validation dataset. The amount of components was selected with the PRESS statistic, which was calculated separately for the predicted size and shape variables. Prior to training and validation, an outlier analysis was conducted on the Kendall tangent space coordinates with function hclust using Ward’s method, a hierarchical clustering method. In this cluster analysis, the Euclidean distance was used as a distance measure between subjects.

### Test dataset and error measure

In order to select a test dataset, a stratified random sample was carried out using function stratsample from Version 3.33.2 of software package survey^48^. The strata were determined using gender, site, and age category in years. Predictions of real-size configurations were made for the subjects in the test dataset. For each landmark, the error measure was calculated as the Euclidean distance in mm in size-and-shape space for convenience of interpretation and communication. For each error measure variable, the statistical expectation was calculated using the log-normal distribution. The resulting expectation variable was transformed to its natural logarithm and clustered using a Gaussian finite mixture model, which was performed with function Mclust of Version 5.4.1 of software package mclust^49^. To test the amount of clusters and whether clusters have the same variance, the Bayesian Information Criterion (BIC) was used.

Prior to the error measure calculations, the natural logarithm of centroid size and Kendall tangent space coordinates of the test dataset were detracted from noise. This was done with functions estim_sigma and adashrink of Version 1.0 of software package denoiseR^[50]^, performing an adaptive shrinkage of singular values. The noise-detracted Kendall tangent space coordinates were back transformed using equation (6) with cos(*ρ*).

### Benchmark prediction method

The proposed method was benchmarked against a method based on that of Xi, Guo, and Shu26. The same predictor variables were used: gender, occupation, education, number of children, and marital status. In this benchmark study, these variables were augmented by the natural logarithms of configuration height in mm, weight in kg, and age in years. The predicted variables were included in a principal component analysis using function prcomp. Each resulting score variable was modelled separately using a regression tree with function rpart of Version 4.1.13 of software package rpart51. Each regression tree was pruned using function prune after cross validations were performed on the training and validation dataset.

## Results

### Data selection

To insure prediction accuracy, 3827 out of 4431 subjects were selected based on two criteria. They should have complete predictor data and configurations that are not outliers. The majority of outliers were asymmetric configurations with an Euclidean distance between the shape and its reflection above 0.06. The remaining outliers had misplaced surface sliding semilandmarks. Furthermore, 696 of the 2074 landmarks were selected to reduce the amount of variables while predicting as many body characteristics as possible. The selected subjects were divided into a training and validation dataset of 3043 subjects and test dataset of 784 subjects.

### Data description

The composition of the training and validation dataset should be similar to the composition of the test dataset. This similarity can be attained from the distribution of gender, site (Table 2), and predictors (Table 3). In the training and validation dataset, there were 1541 female subjects (50.641%) and 1502 male subjects (49.359%), and in the test dataset 403 female subjects (51.403%) and 381 male subjects (48.597%).

**Table 2.**
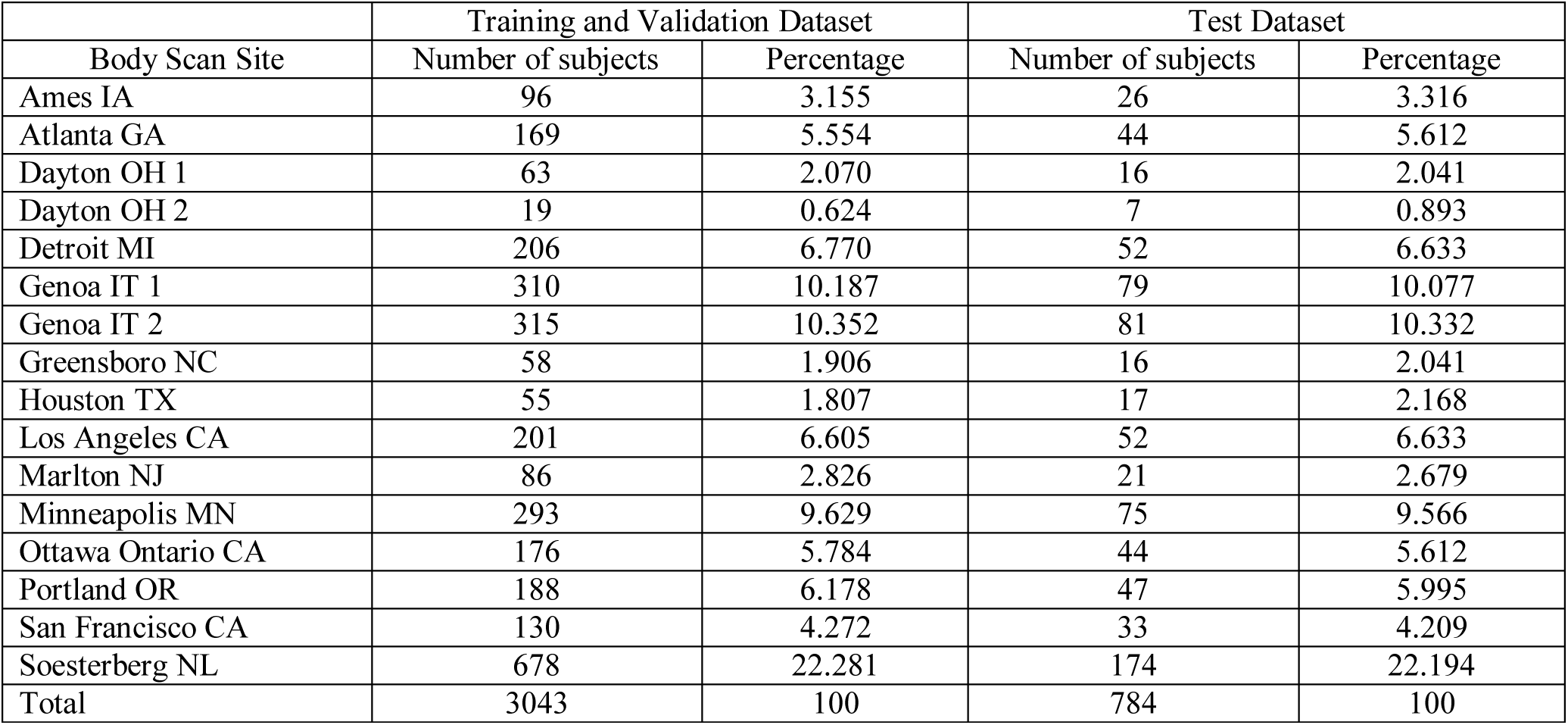
Number of subjects and percentage in the training and validation dataset and test dataset by body scan site. Every name of a body scan site consists of a place name and a state or country abbreviation. The sites are located in North America, Italy, and the Netherlands.

Table 3 shows a group of diverse subjects of which the mean shape has more Caucasian features (Fig. 1). It should be noted that children and adolescents were barely represented.

**Figure 1.**
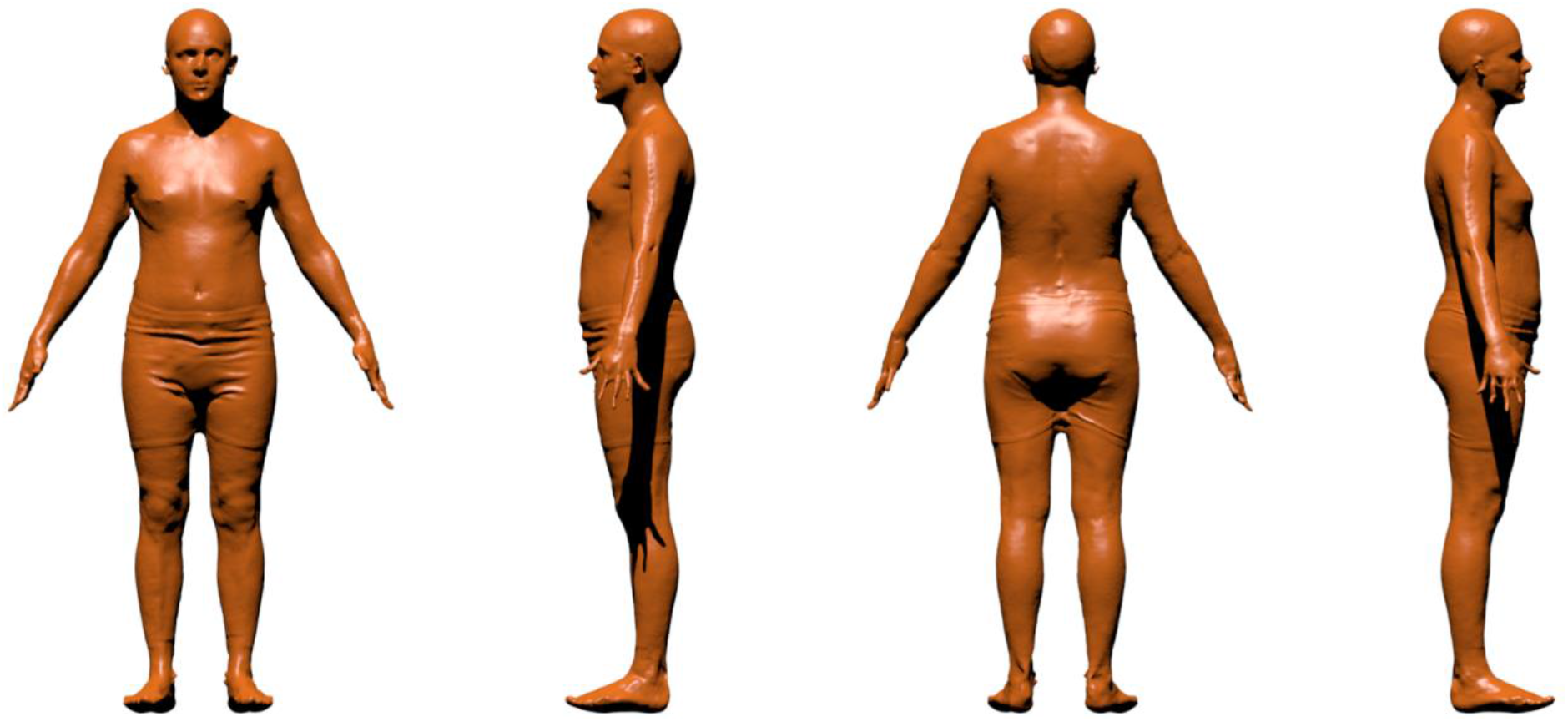
Mesh warped into the mean shape using the thin-plate spline method. The selected mesh is from a subject (made unrecognisable due to the warping) whose shape is closest to the mean shape.

**Table 3.** Statistics of predictor variables in the training and validation dataset and test dataset. The expectation and standard deviation were calculated using the log-normal distribution.

| Variable | Statistic | Training and Validation Dataset | Test Dataset |
| --- | --- | --- | --- |
| Configuration height in mm | Minimum | 1403.995 | 1491.866 |
|  | Maximum | 2055.320 | 2035.530 |
|  | Expectation | 1726.121 | 1728.641 |
|  | Standard deviation | 95.594 | 100.613 |
| Weight in kg | Minimum | 37.900 | 43.400 |
|  | Maximum | 156.690 | 154.880 |
|  | Expectation | 73.679 | 74.284 |
|  | Standard deviation | 16.382 | 17.587 |
| Age in years | Minimum | 17.000 | 18.000 |
|  | Maximum | 68.000 | 65.000 |
|  | Expectation | 36.878 | 37.230 |
|  | Standard deviation | 13.540 | 13.933 |

### Model results

The orthogonal projection, kernel partial least squares regression model, and principal component analysis performed well. With regard to the orthogonal projection, the results are a squared Pearson correlation (r^2^) of 0.999731 and regression equation intercept (α) of 0.000436 and slope coefficient (β) of 0.984494. The kernel partial least squares regression model was tested with 80 components and a 5-fold cross-validation. For size, represented by the natural logarithim of centroid size, 0 component gave a maximum PRESS statistic of 9.430735, and 43 components showed a minimum PRESS statistic of 0.196990. For shape, represented by Kendall tangent space coordinates, 0 component gave a maximum mean PRESS statistic of 0.000619, and 39 components indicated a minimum mean PRESS statistic of 0.000433. A total of 39 components were chosen because more components did not significantly reduce the size PRESS statistic. With regard to the principal component analysis, the first 10 principal components were used and explained approximately 95.428% of the total variance.

### Prediction results

In the test dataset, surface sliding semilandmarks were more likely to be accurately predicted than fixed landmarks. For every landmark the mean error was calculated, and the resulting means were combined into one expectation variable which was transformed to its natural logarithm and clustered. To find the amount of clusters and variance specification per cluster that best fit the data, the mixture model was calculated for equal and unequal variance between clusters, for 1 up to and including 10 groups.

With regard to the proposed method, for unequal variance and 2 groups the BIC was 76.792 and optimal. For Cluster 1 the *a-priori* probability coefficient, expectation, and standard deviation of the log-normal distribution were 0.824, 8.816, and 1.180; for Cluster 2, these parameters were 0.176, 18.454, and 8.844. Cluster 1 consisted only out of 597 surface semilandmarks (100.000%), and cluster 2 out of 25 surface semilandmarks (25.253%) and 74 fixed landmarks (74.747%).

As for the benchmark method, for unequal variance and 2 groups the BIC was 51.017 and optimal. For Cluster 1 the *a-priori* probability coefficient, expectation, and standard deviation of the log-normal distribution were 0.827, 8.702, and 1.182; for Cluster 2, these parameters were 0.173, 19.030, and 8.950. Cluster 1 consisted only out of 596 surface semilandmarks (100.000%), and cluster 2 out of 26 surface semilandmarks (26.000%) and 74 fixed landmarks (74.000%).

## Discussion

Based on hormonal literature, cells in the whole body can respond to hormones. This means that if the response of some of the cells is known, the response of others can be predicted. This can be evidenced by the prediction results of the method proposed in this manuscript.

The expected error of landmarks was mostly below 10 mm, making the hormonal framework useful in practical prediction applications. For example. in online fashion retail approximately 10 mm and 40 mm can be the minimum and maximum size difference of clothing. Thus, from a practical perspective, the prediction method performs well.

With regard to scientific contribution, the author’s proposed prediction method outperforms existing methods in at least two crucial aspects. The method of Zhu and Mok^25^ had an average of mean absolute size discrepancies of six different girth measurements of 9.690 mm in the case of loose-fit clothing, but the following should be taken into account. On the one hand, Zhu and Mok tested their method with only 21 subjects (15 female and 6 male), while the current method was tested with 784 subjects. On the other hand, Zhu and Mok required two photos of the whole dressed body, while the current method only required a passport photo. The benchmark method in this paper also will require more input in the form of subjective answers.

Besides prediction accuracy, the proposed prediction method has several strengths and one weakness. The first and foremost strength is that the method requires little input from respondents: only a passport photo and answers to three closed-ended questions. Second, the method has a theoretical foundation of hormones, making it easier to improve the prediction method in the future. For example, hormones can be related to behaviour, and thus the prediction of size and shape of the human body can be enhanced by using answers to questions about behaviour as predictors. Third, one statistical model (rather than multiple) is used in the method, reducing accumulated prediction error. Fourth, the prediction results can be visualised using the thin-plate spline method. A weakness of the prediction method is that not all hormonal responses of the body can be predicted with the shape of the face. For example, the face shape does not respond to dihydrotestosterone, which is crucial in development of the penis^12^.

The prediction method was tested with the CAESAR dataset, which had several limitations that can be opportunities for future research. First, the dataset contained predominantly Caucasian subjects and virtually no children and adolescents, and the number of subjects was too small to incorporate many nuances between subjects. Future investigation should include more diverse populations and subjects. Second, the body posture could be inconsistent across subjects, making configurations asymmetric at times, and therefore more difficult to predict. Future research would benefit by using a 3D body scan dataset in which subjects strictly adhere to a detailed posture protocol. The CAESAR project included such a protocol^27^, but wasn’t very thorough, which allowed subjects some freedom of physical movement. Of course, future research could also use posture correction methods before testing the prediction method, but these algorithms may add reconstruction error to the prediction error. Third, fixed landmarks coordinates probably contained landmark placement error. This is because prediction results showed that fixed landmarks were harder to predict than surface semilandmarks, which were corrected by sliding with the Procrustes distance^34^. Future research could benefit by creating a 3D body scan dataset with multiple body scans and placements of all fixed landmarks per subject. This makes it possible to calculate, check, and correct the landmark placement error per subject. Fourth, the amount of details in shape and color of the body scans were low, especially in the dataset of the Netherlands. Future research would benefit from body scan datasets using more sophisticated body scan technology (the CAESAR dataset dates from 2002).

## Acknowledgements

The author is grateful to Dr. Cristina Tortora of the San José State University for helpful comments that have improved the geometric morphometrics in this article. This work was funded by Krossfields.

## Author contributions

J. v. V. is the sole contributor to this manuscript and the prediction method therein.

## Data availability

The CAESAR dataset can be retrieved from https://wear.wildapricot.org.

